# Brain simulation augments machine-learning-based classification of dementia

**DOI:** 10.1101/2021.02.27.433161

**Authors:** Paul Triebkorn, Leon Stefanovski, Kiret Dhindsa, Margarita-Arimatea Diaz-Cortes, Patrik Bey, Konstantin Bülau, Roopa Pai, Andreas Spiegler, Ana Solodkin, Viktor Jirsa, Anthony Randal McIntosh, for the Alzheimer’s Disease Neuroimaging Initiative, Petra Ritter

**Affiliations:** Berlin Institute of Health at Charité – Universitätsmedizin Berlin, Berlin, Germany; Department of Neurology with Experimental Neurology, Brain Simulation Section, Charité – Universitätsmedizin Berlin, corporate member of Freie Universität Berlin and Humboldt-Universität zu Berlin, Berlin, Germany; Institut de Neurosciences des Systèmes, Aix Marseille Université, Marseille, France; Bernstein Center for Computational Neuroscience Berlin, Berlin, Germany; Department of Neurophysiology and Pathophysiology, University Medical Center Hamburg-Eppendorf, Hamburg, Germany; Neuroscience, Behavioral and Brain Sciences, UT Dallas. Richardson, TX, USA; Baycrest Health Sciences, Rotman Research Institute, Toronto, Ontario, Canada

**Author notes:** Corresponding author: Prof. Dr. med. Petra Ritter, Berlin Institute of Health, Department of Neurology with Experimental Neurology, Brain Simulation Section, Robert-Koch-Platz 4, 10115 Berlin, Germany. The authors Paul Triebkorn and Leon Stefanovski had equal contributions to this article. Data used in preparation of this article were obtained from the Alzheimer’s Disease Neuroimaging Initiative (ADNI) database (adni.loni.usc.edu). As such, the investigators within the ADNI contributed to the design and implementation of ADNI and/or provided data but did not participate in analysis or writing of this report. A complete listing of ADNI investigators can be found at: http://adni.loni.usc.edu/wp-content/uploads/how_to_apply/ADNI_Acknowledgement_List.pdf.

**Keywords:** Alzheimer’s Disease, The Virtual Brain, Multi-scale brain simulation, Machine Learning, Positron Emission Tomography

## Abstract

**INTRODUCTION:** Computational brain network modeling using The Virtual Brain (TVB) simulation platform acts synergistically with machine learning and multi-modal neuroimaging to reveal mechanisms and improve diagnostics in Alzheimer’s disease.

**METHODS:** We enhance large-scale whole-brain simulation in TVB with a cause-and-effect model linking local Amyloid β PET with altered excitability. We use PET and MRI data from 33 participants of Alzheimer’s Disease Neuroimaging Initiative (ADNI3) combined with frequency compositions of TVB-simulated local field potentials (LFP) for machine-learning classification.

**RESULTS:** The combination of empirical neuroimaging features and simulated LFPs significantly outperformed the classification accuracy of empirical data alone by about 10% (weighted F1-score empirical 64.34% vs. combined 74.28%). Informative features showed high biological plausibility regarding the Alzheimer’s-typical spatial distribution.

**DISCUSSION:** The cause-and-effect implementation of local hyperexcitation caused by Amyloid β can improve the machine-learning-driven classification of Alzheimer’s and demonstrates TVB’s ability to decode information in empirical data employing connectivity-based brain simulation.

**RESEARCH IN CONTEXT:**

1. **SYSTEMATIC REVIEW**. Machine-learning has been proven to augment diagnostics of dementia in several ways. Imaging-based approaches enable early diagnostic predictions. However, individual projections of long-term outcome as well as differential diagnosis remain difficult, as the mechanisms behind the used classifying features often remain unclear. Mechanistic whole-brain models in synergy with powerful machine learning aim to close this gap.
2. **INTERPRETATION**. Our work demonstrates that multi-scale brain simulations considering Amyloid β distributions and cause-and-effect regulatory cascades reveal hidden electrophysiological processes that are not readily accessible through measurements in humans. We demonstrate that these simulation-inferred features hold the potential to improve diagnostic classification of Alzheimer’s disease.
3. **FUTURE DIRECTIONS**. The simulation-based classification model needs to be tested for clinical usability in a larger cohort with an independent test set, either with another imaging database or a prospective study to assess its capability for long-term disease trajectories.

## 1. INTRODUCTION

Although the spectrum of Alzheimer’s Disease (AD^†^)-related disease burden is broad and its early diagnosis a common modern health problem, the knowledge of underlying disease mechanisms remains incomplete. Besides the two hallmark proteins Amyloid β (Aβ) [1, 2] and Tau [3, 4], other involved factors have been identified, such as e.g., impairment of the blood-brain-barrier [5], synaptic dysfunction [6], network disruption [7], mitochondrial dysfunction [8], neuroinflammation [9] as well as genetic risk factors [10]. While Aβ and Tau are widely accepted as involved core features [11, 12], their mutual interaction [13] and interaction with other factors [5] are incompletely understood. Comprehensive knowledge of this multifactorial interaction in the pathogenesis of AD is crucial for further therapeutic strategies, including recent developments of potentially disease-modifying anti-Aβ therapy with Aducanumab [14].

The Virtual Brain (TVB, www.thevirtualbrain.org) is an open-source platform for modeling and simulating large-scale brain networks by using personalized structural connectivity models [15, 16]. TVB enables the model-based inference of underlying neurophysiological mechanisms across different brain scales that are involved in the generation of macroscopic neuroimaging signals including functional magnetic resonance imaging, electroencephalography (EEG) and magnetoencephalography. Moreover, TVB facilitates the reproduction and evaluation of individual configurations of the brain by using subject-specific data. In this study, we make use of virtual local field potentials (LFPs) from simulated brain data from a recent experiment with TVB [17]. We integrate individual Aβ patterns, obtained from positron emission tomography (PET) with the Aβ-binding tracer ^18^F-AV-45, into the brain model. Consecutively, distinct spectral patterns in simulated LFPs and EEG could be observed for patients with AD, mild cognitive impairment (MCI), and healthy control (HC) subjects (**Figure 1**). Such integration was done by transferring the local concentration of Aβ to a variation in the brain model’s local excitation-inhibition balance. This resulted in a shift from alpha to theta rhythms in AD patients, which was located in a similar pattern as local hyperexcitation in core structures of the brain network. The frequency shift was reversible by applying “virtual memantine”, i.e., virtual N-methyl-D-aspartate (NMDA) antagonistic drug therapy. An overview of the study results is provided in **Figure 1**.

**Figure 1.**
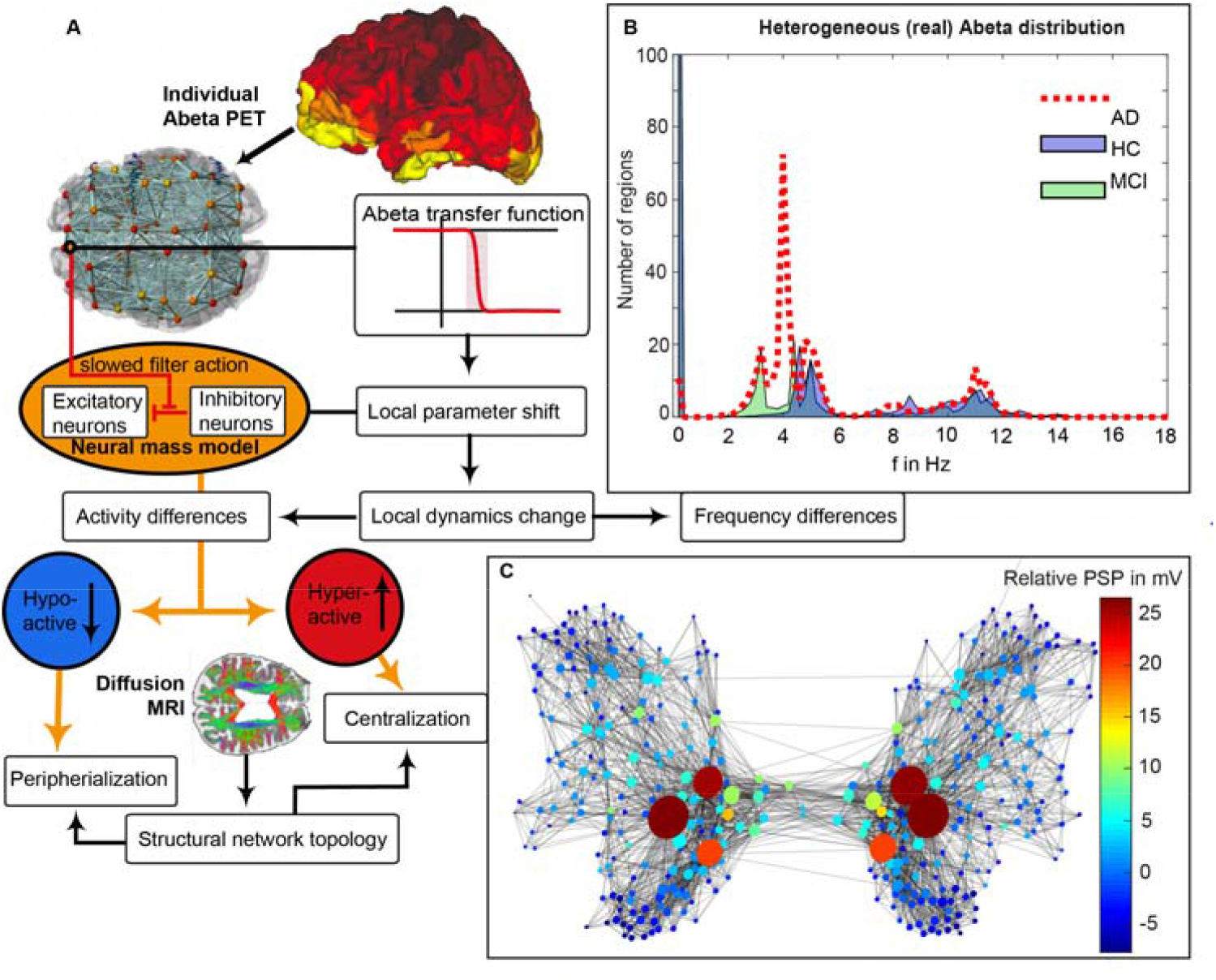
Modified from [17]. AβPET-driven brain simulation model of AD. (A): Regional PET intensity constraints regional parameters. A sigmoidal transfer function translates the regional Aβ load to changes in the excitation-inhibition balance. (B) Virtual AD patient brains exhibited significantly slower simulated LFPs than MCI and HC virtual brains and showed a shift from alpha to theta frequency range. While the AD group is solely dominated by two clusters in the alpha and theta band, the groups of HC and MCI have an additional strong cluster exhibiting no oscillations (“frequency of zero”), called a stable focus. This phenomenon is absent in the AD group. The stable focus in HC and MCI virtual brains provides an additional – simulation inferred - distinctive criterion between groups. The graph in (C) represents the SC, wherein the nodes’ size reflects the degree, while color corresponds to the relative postsynaptic potentials (relative to the mean postsynaptic potential of the simulation). The graph indicates that local hyperexcitation occurs in central parts of the networks. Moreover, the observed slowing was spatially associated with local hyperexcitation.

AD-specific pathologies, such as deposition of Aβ in neuritic plaques, Tau deposition in neurofibrillary tangles, and atrophy of neural tissue, have been widely studied with machine learning (ML) approaches [18, 19]. The major advantage of employing ML-based classification algorithms on neuroimaging data is the potential for recognizing complex high dimensional, previously unknown disease patterns in the data, potentially identifying AD before clinical manifestation or predicting a disease trajectory.

In this research, we hypothesize that TVB-inferred features improve the classification performance compared to imaging data features alone.

We further argue that the current sample size of 33 subjects is sufficient to achieve a reliable proof-of-concept, considering the following three main aspects:

1. This study aims to show an information gain provided by TVB with regards to differential classification between HC, MCI and AD populations. We do not aim to push generalizability performance of state-of-the-art machine learning methodologies with this sample size. This leads to a primary focus on the group level significance of the decoding accuracy rather than the accuracies themselves [20].
2. This information gain and the significance of the model performances are validated by comparing the distributions of model accuracies between feature sets and against null distributions of accuracies approximated using permutation testing [20].
3. As implemented in our approach, nested cross-validation still represents the best way to estimate generalizability in the given context [21]. In combination with the previous points, this leads to a feasible and robust estimation of the information gain.

We show that TVB simulations provide additional unique diagnostic information that is not readily available using the available empirical data alone. This supports the idea that TVB provides value and real-world applicability above and beyond merely reorganizing empirical data, and suggests that its simulations are biologically plausible.

## 2. MATERIALS AND METHODS

### 2.1. Alzheimer’s Disease Neuroimaging Initiative (ADNI) database

Data used in the preparation of this article were obtained from the Alzheimer’s Disease Neuroimaging Initiative (ADNI) database (adni.loni.usc.edu). The ADNI was launched in 2003 as a public-private partnership, led by Principal Investigator Michael W. Weiner, MD. The primary goal of ADNI has been to test whether serial magnetic resonance imaging (MRI), PET, other biological markers, and clinical and neuropsychological assessment can be combined to measure the progression of MCI and early AD. For up-to-date information, see www.adni-info.org.

### 2.2. Data acquisition, processing, and brain simulation

The detailed methodology of data acquisition, selection, processing, and simulation are described in a former study [17]. We included 33 ADNI-3 participants, thereof 10 AD patients, 15 HC participants and eight MCI patients. The selection criteria included availability of both Aβ and Tau PET, diffusion-weighted MRI, and all MRI sequences necessary to fulfill the standards of the human connectome project minimal preprocessing pipeline [22].

In addition to the data used in our former study [17], we also used the distribution of Tau in ^18^F-AV-14-51 PET for our analyses to obtain the best available empirical data basis. Again, the nuclear signal intensity is related to a reference volume in the cerebellum.

For the subcortical volumetrics used in this study, we obtained the volumetry statistics provided by the -*autorecon2* command. The segmentation is performed with the modified Fischl parcellation [23] of subcortical regions in Freesurfer [24].

A detailed description of image processing can be found in **Appendix A**.

Whole-brain simulations with TVB are based on a structural connectivity (SC) matrix derived from diffusion-weighted MRI. After processing the empirical imaging data, we used the SC of the HC population to generate an averaged standard SC for all participants. For the simulations, we made use of the Jansen-Rit neural mass model [25, 26]. Neural mass models employ a mean field simplification to compute electrical potentials on a regional level by using oscillatory equations systems [27]. The variables, parameters and model equations can be found in [17]. Parameter settings were chosen due to theoretical considerations in former studies[17, 28]. We explored a range of the global scaling factor G, a coefficient that scales the connection between distant brain regions, to capture different dynamic states of the simulation. The novelty in our recent simulation study was the introduction of a mechanistic model for Aβ-driven effects. We linked local Aβ concentrations, measured by Aβ PET, to the excitation-inhibition balance in the model by defining the inhibitory time constant τ_i_ as a sigmoidal function of local Aβ burden [17].

The simulation models electrical potentials in the whole brain, here measured on the region level by LFPs. In addition, we calculate the EEG signal as a projection of the LFP from within the brain to the surface of the head, taking into the concept of a lead-field matrix simplification to three compartment borders brain-skull, skull-scalp, and scalp-air [15, 29-31].

A detailed description of the simulations can be found in **Appendix B**.

### 2.3. Machine Learning Approach

Our primary objective is to determine if extracted features from TVB add to the classifiers’ predictive power. To achieve this, we repeated the ML procedure with three different feature sets: a) using empirical features alone, b) using simulated features alone, and c) combining both types of features into a hybrid model.

As empirical features, we employed 380 regional values for each Aβ PET standardized uptake value ratio (SUVR) and Tau PET SUVR, and 40 subcortical volumes, leading to 800 empirical features. As simulated features, we used 379 regional LFP frequencies from the simulations. The combined feature space contains all the above with 1179 features.

Two types of ML classifiers that are suitable for small-sample classification problems were used: the kernel-based support vector machines (SVM) [32] and the decision-tree based Random Forest (RF) [33].

By training two classifiers based on different underlying machine learning mechanisms, we provide more robust evidence that the pattern in classification performance, when combining simulated and empirical features, is reliable and clinically relevant. Further, this pattern is driven by a reliably reoccurring subset of the features themselves, rather than by particular mechanisms underlying a classification algorithm.

Our ML approach is designed to fulfill primarily two goals:

1. Providing a robust, reproducible, and accurate evaluation of classification performance with the data.
2. Facilitating exploration of the empirical and simulated features that are most important for achieving optimal separation between the AD, MCI, and HC groups.

To satisfy the first goal, we implement a strict nested cross-validation scheme that allows us to obtain statistically reliable classification performance metrics while minimizing overfitting in a P >> N setting (i.e., we have a small sample size N, but a very large number of features P). Our cross-validation method is adapted from earlier work in machine learning for clinical neuroscience [34], and is described in greater detail in **Figure 2**.

**Figure 2.**
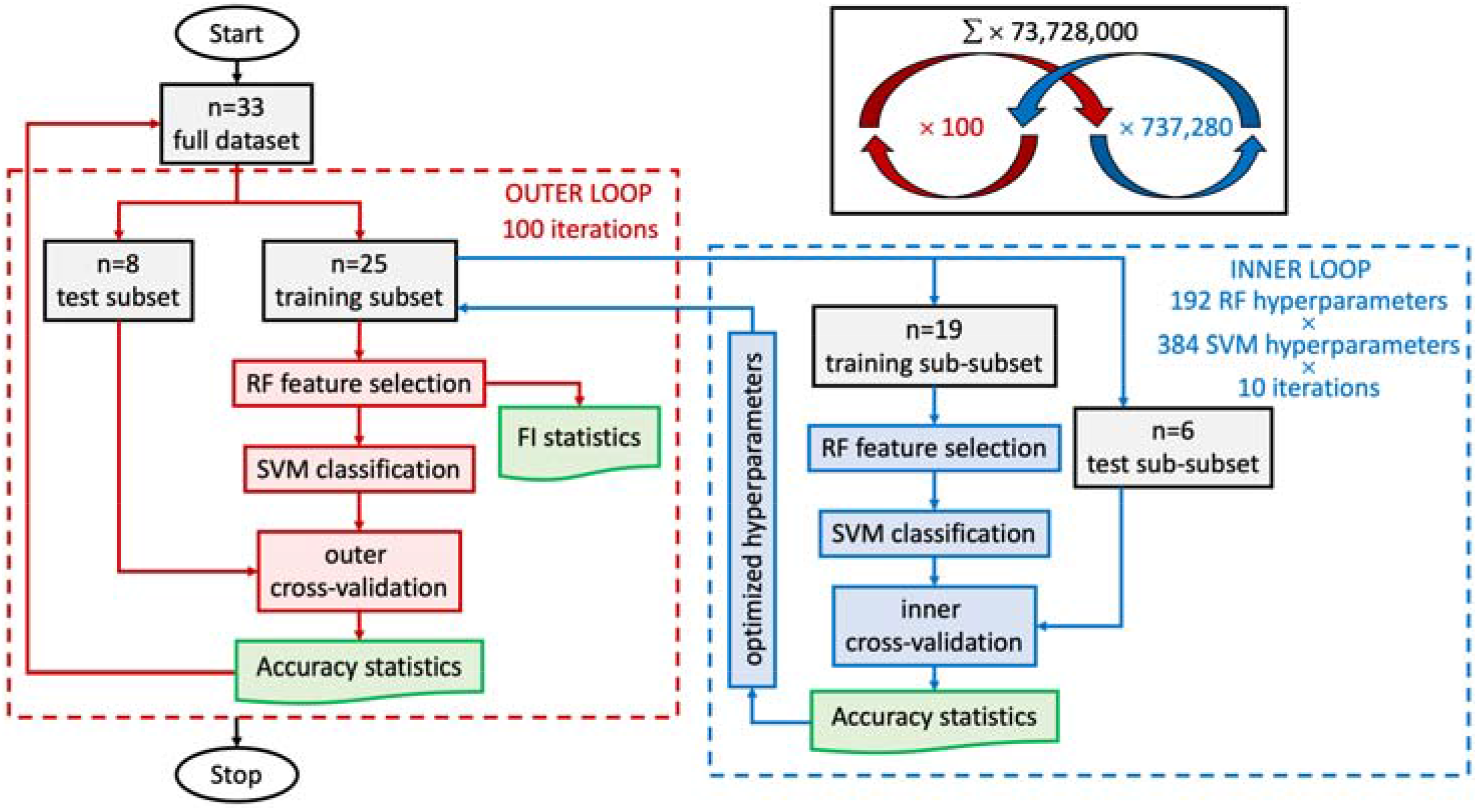
Nested cross-validation loop design. Starting in the outer loop: stochastic cross-validation starts with 100 iterations using 25% of data (randomly selected per iteration) for testing. The training subset goes to the inner loop after the train-test split. In the inner loop: split data again just like in the outer loop to obtain training set and validation set for an inner 10 cross-validation iterations with each hyperparameter setting (in total 192 combinations for RF and 384 for SVM, leading to 73,728 combinations with every 10 iterations). Next, we scale training features by subtracting the median and dividing by the inter-quartile range (makes them robust to outliers we identified above). We apply these scaling statistics calculated from the training set also to the test set. Then, we iterate through hyperparameters (**Supplementary Tables C.1 and C.2**). RF is used for feature selection. Afterward, the remaining features are used for training the SVM classifier with specific hyperparameter settings. We track the selected features for each run and compute the frequency with which they are selected across iterations for the outer loop. The SVM classifications are validated with the test sub-subset (inner cross-validation). This provides optimized hyperparameter settings from the inner cross-validation loop. Back to the outer loop, we recombine training and validation data (which were separated in the inner loop) - still keeping test data separate. We set hyperparameters to the best settings obtained in the inner loop. Then, we train the model and record results: RF is again used for feature selection, which leads to feature importance (FI) statistics used for the results. Afterward, SVM classifies the reaming features, which are then validated with the test set (outer cross-validation). After this, the next iteration of the outer loop begins.

We satisfy the second goal in two ways. First, our cross-validation scheme provides a natural metric for feature relevance, i.e., feature selection frequency across cross-validation runs. Additionally, we use feature importance metrics inherent to each feature selection method explored. In our case, the F-statistic and the entropy criterion were two metrics used for feature selection for the SVM and the RF respectively.

Currently, the most reliable method for statistical control of prediction accuracy is permutation testing [20]. To this end, we performed the same classification pipeline, including all feature preprocessing, feature selection and cross-validation steps, using randomly shuffled class labels. This was repeated 750 times to achieve a robust estimate of the null model as an approximation for the inherent prediction error of the model and chance classification results.

A detailed technical description of the ML methodology can be found in **Appendix C**.

## 3. RESULTS

### 3.1. Data properties

We used basic descriptive statistics to assess data quality prior to machine learning analysis. The distribution of simulated LFP frequencies, Aβ PET SUVR, Tau SUVR, and regional Volumes and their interdependency are shown in **Figure 3**. Aβ (*P* = .0018) and Tau SUVR (*P* = .0005) are significantly different between AD and HC after Bonferroni-correction. LFP frequency differs significantly between AD and MCI (*P* = .0316) but is not significant after Bonferroni-correction. Interestingly, in the existing data, there is no significant difference in the mean volumes between groups.

**Figure 3.**
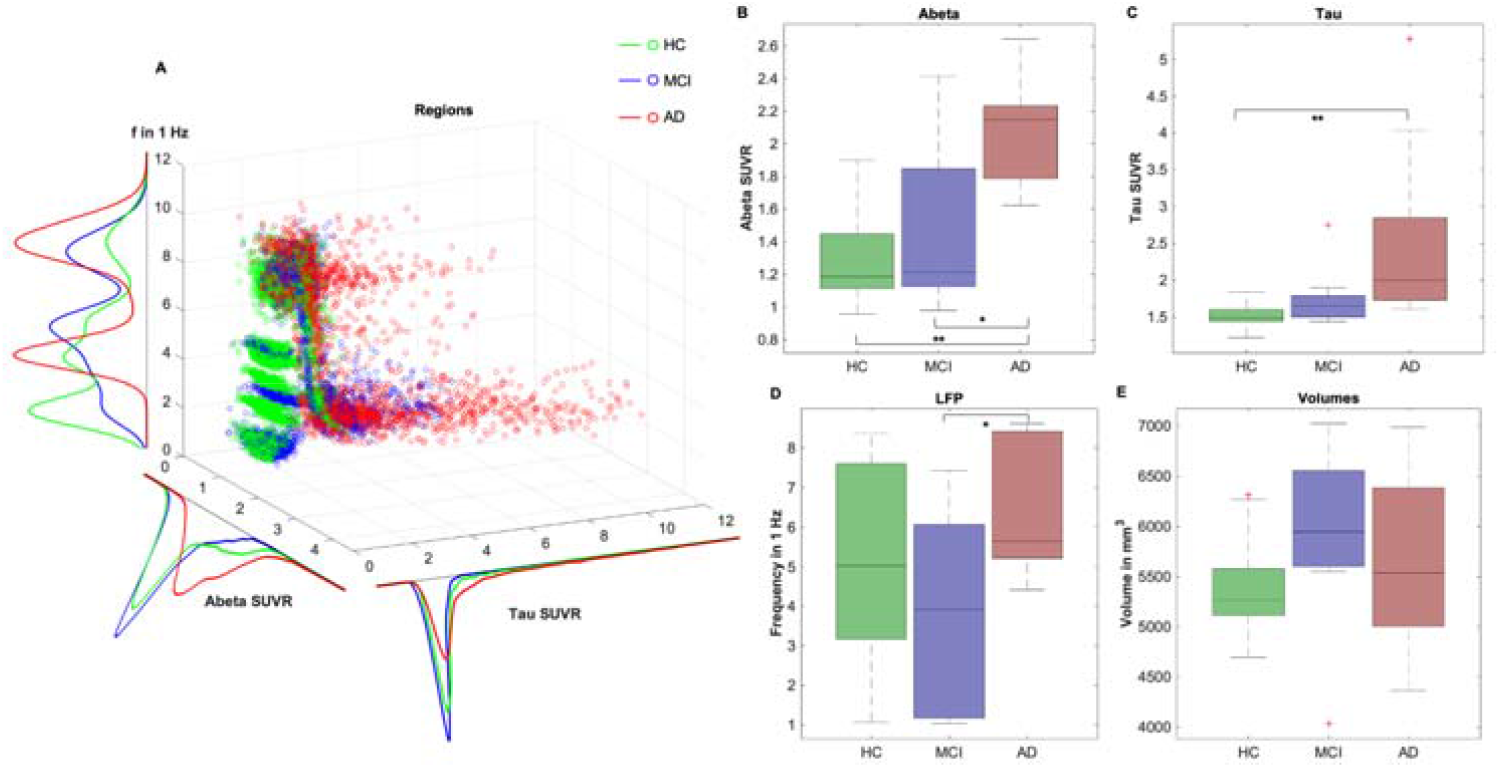
Characteristics of empirical feature space. In (**A**), regional distributions of Aβ, Tau, and LFP frequency are shown for all groups in a 3D scatterplot. Red datapoints symbolize regions of AD patients, green points MCI patients, and blue points HC. Each scatters point stands for one region of one subject. Color density is normalized between groups. A kernel density estimate of the corresponding histograms is shown (projection of the 3D-plot to one axis). In particular, it can be seen a string of outliers with very high Tau values in the AD group and in parts in the MCI group, which does not appear for HC. Moreover, AD participant’s regions show higher Aβ values, in particular for lower frequencies. Besides, boxplots are presented for groupwise comparisons for the features mean Aβ per subject, mean Tau per subject, mean simulated LFP frequency per subject, and mean Volume per subject. A Kruskal-Wallis-test was performed to assess significance: * marks significance with *P* < .05; ** marks significance after Bonferroni-correction with *P* < .0042. (**B**) Aβ SUVR is significantly different between AD and HC (*P* = .0018) and MCI (*P* = .0454), but not between HC and MCI (*P* = .8113). (C) Tau SUVR is only significantly different between AD and HC (*P* = .0005), but not between AD and MCI (*P* = .1736) or HC and MCI (*P* = .2671). (D) LFP frequency is only significantly different between AD and MCI (*P* = .0316), but not between AD and HC (*P* = .2160) or HC and MCI (*P* = .4716). (E) The mean Volume of all regions does not show significant differences: AD and MCI (*P* = .7056, AD and HC (*P* = .5102) or HC and MCI (*P* = .1405).

### 3.2. Classification performance

Overall, we performed nine experiments with three different classification schemes and three feature sets (see **Appendix D**). The hybrid classification scheme with SVM and RF performed best. For all schemes, the combined feature space outperformed both the empirical and the simulated feature space (**Supplementary Table D.1**).

In the hybrid classifier scheme, RF was used in the inner cross-validation loop (10 iterations) to perform multivariate feature selection. In the outer cross-validation loop (100 iterations), the features used by the best random forest model were then used to train an SVM.

Weighted F1-score and normalized confusion matrices are given in **Figure 4**. The combined approach (wF1 = 0.7428) outperformed the empirical one (wF1 = 0.6434) by about 0.1 (**Figure 4D**), mainly because of an improvement in the classification of the MCI group (**Figure 4A-C**). We used the Wilcoxon Signed-Rank test from 100 cross-validation runs to assess significance (Shapiro-Wilk test of normality for the F1 score distributions revealed *P* < .01 for empirical and combined approach and *P* = .07 for the simulated approach, leading to the usage of a non-parametric test). The differences between the combined approach and both other approaches were highly significant (combined and empirical: *P* = 2.52 ×10^−11^; combined and simulated: *P* = 1.6 ×10^−7^), meanwhile there was no significant difference between the empirical and simulated approaches (*P* = .34).

**Figure 4.**
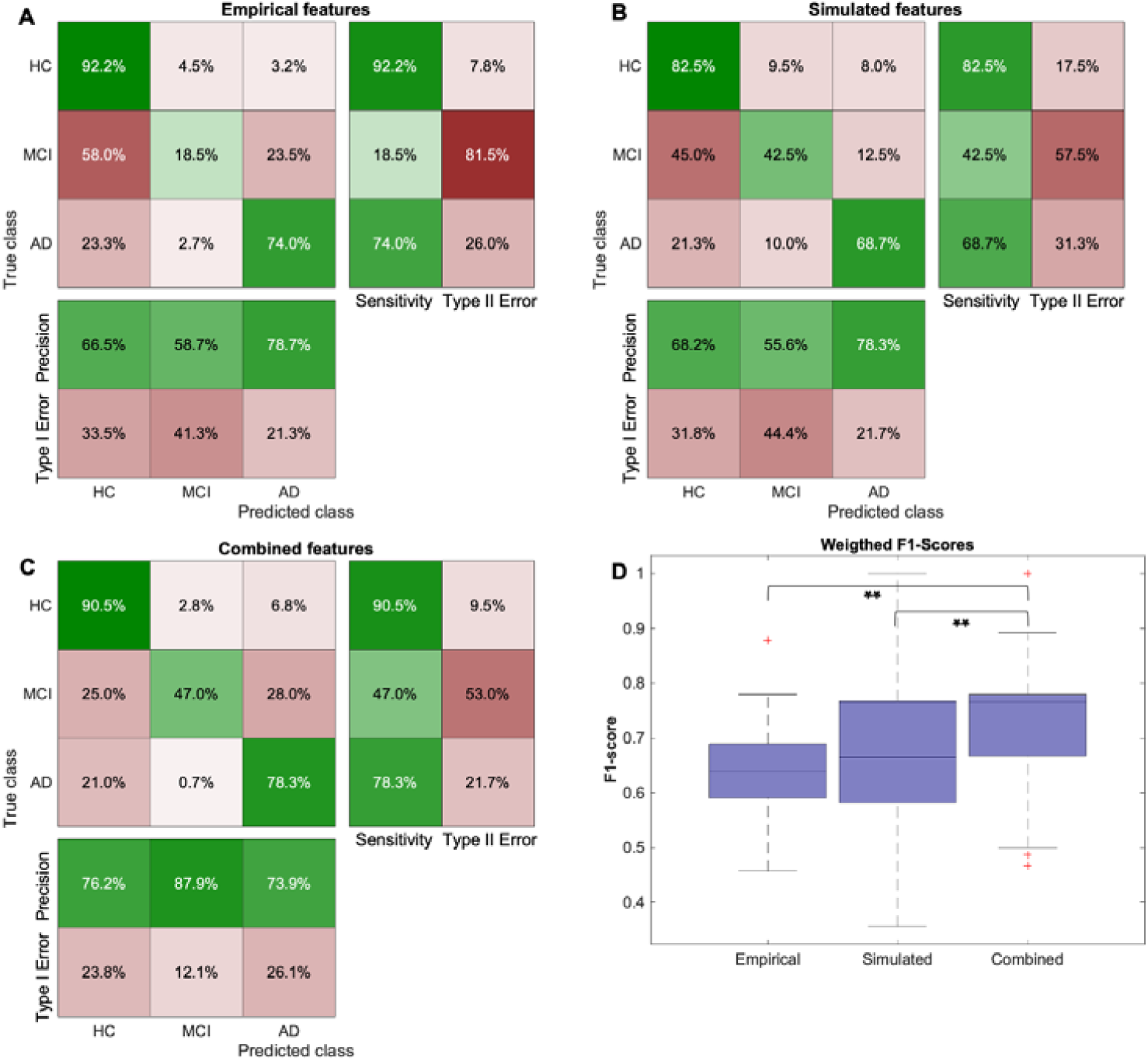
Results of the nested cross-validation classification approach. (**A-C**) Confusion matrices are computed by summing the confusion matrices across all 100 cross-validation runs and normalizing per class. In particular, the combined approach improved the prediction of MCI participants, as AD and HC were already quite well distinguishable by the empirical features. (**D**) Boxplots of mean weighted F1-scores for three different feature spaces. The combined approach (0.7428) outperformed the empirical one (0.6434) by about 0.1. Significance assessment with the Wilcoxon Signed-Rank test from 100 cross-validation runs: combined vs. empirical: *P* = 2.52 ×10^−11^; combined vs. simulated: *P* = 1.6 ×10^−7^, empirical vs. simulated *P* = .34.

### 3.3. Classification validity

As a further analysis to understand this classification improvement, we calculated the feature importance. **Figure 5A** shows the mean entropy-based feature importance given by the Random Forest classifier for 100 outer cross-validation runs. This is used to show that there is a decreasing curve, as we would expect if meaningful features were found (as opposed to a more uniform distribution). Many of the more important features seem to be biologically plausible in the context of AD (**Figure 5B**), as, e.g., Tau in entorhinal cortex (Braak stage 1), frequencies in the thalamus (as significant rhythm generator) and putamen, and volumes in putamen and hippocampus (as signs of atrophy). A table with the full name and corresponding functional network according to Rosen and Halgren [35] for the 50 highest ranked features is given in **Supplementary Table D.2**.

**Figure 5.**
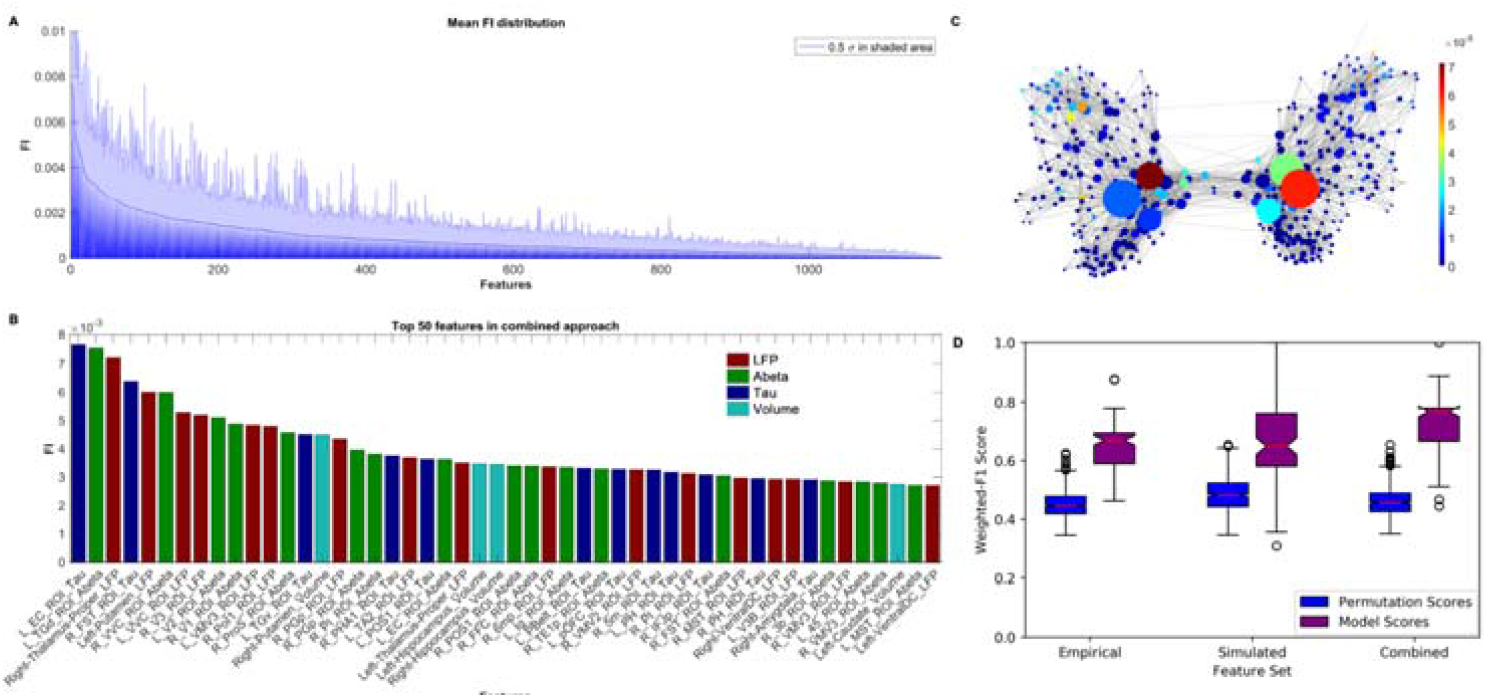
Feature importance (FI) distribution. (**A**) Mean RF-derived feature importance from 100 outer cross-validation runs. Entropy criterion with combined feature types shown here. Feature importance values are normalized, so all features sum to 1. In shaded blue, half standard deviation is displayed for each feature. (**B**) Top 50 features across all cross-validation runs. Both empirical (Tau in dark blue, Aβ in green, Volume in light blue), as well as simulated frequencies (red), contributed to the improved classification. Many features seem moreover to be biologically plausible in the context of AD, e.g., Tau in the entorhinal cortex, thalamic and putaminal frequencies, and putaminal as well as hippocampal volumes. (**C**) Visualization of the SC graph with color indicating FI of the regional LFP frequencies, while vertex diameter reflects the structural degree. It shows a network dependency of the LFP FI. Only edges with connection strength above the 95^th^ percentile are shown. (**D**) The distributions of weighted F1 scores for permutation based null model (left box) and corresponding true model (right box). All models significantly outperform the null model with the combined model showing the greatest average distance to its null model, indicating the gain in differentiating information.

We also showed that feature relevance is dependent on the structural degree of the regions in the underlying SC network (**Figure 5C**). This is an indicator of network effects contributing to the improved classification and another indicator for meaningful classification results.

Using the Wilcoxon signed-rank test, we could further show that the classification performance was significantly higher than the null model (with *P* values of 2.2 ×10^−15^ (simulated), 7.8 ×10^−18^ (empirical) and 4.8 ×10^−18^ (combined)). The average performance of the combined approach showing the greatest distance to the corresponding null model laying outside the 100% interval (**Figure 5D**).

## 4. DISCUSSION

In this study, we show that the involvement of virtual, simulated TVB features into ML classification can lead to an improved classification between HC, MCI, and AD.

The diagnostic value of the underlying empirical features can be improved by integrating the features into a multi-scale brain simulation framework in TVB. We showed for ML algorithms a superiority of feature sets that contain both the empirical and the virtual derived metrics. The absolute gain of accuracy was 10 %. Keeping in mind that all differences between the subjects have to be derived from their Aβ PET signal (because all other factors, e.g., the underlying SC, are the same) this provides evidence that TVB is able to decode the information that is contained in empirical data like the amyloid PET. More specific for the PET and its usage in diagnostics, it highlights the relevance of spatial distribution, which is often not considered in its analysis.

The main reason for this improvement seems to be a better classification of MCI subjects. without the simulated features, the models frequently misclassify MCI subjects as HC. In contrast, the simulated features alone result in more misclassification of healthy controls as either MCI or AD subjects compared to using the empirical features alone. However, combining the empirical features with the simulated features appears to complement their strengths in a clinically useful way; these models retain all or most of the ability to correctly classify healthy controls with the empirical features and retain much of the simulated features’ ability to classify MCI patients. The processing inside TVB seems to reorganize the existing data beneficially.

In theory, a larger number of available features could provide a machine learning algorithm greater flexibility in finding useful combinations. This is the case simply due to a higher degree of freedom during feature selection and weighting. However, the equal empirical data foundation (only PET as individual features) in combination with nested cross-validation method protects from an overfitting bias due to the larger feature space, with additional evidence of this provided by the chance level performance of the null distributions. If the explanation for the improvement in classification accuracy was simply the presence of additional noisy features, we would see a flatter feature importance distribution than shown in **Figure 5**, and therefore a more random distribution of selected features across the 100 cross-validation iterations. Instead, we see that only a few features with high importance are consistently guiding classification, indicating that they in fact provide useful discriminative information. Preventing this kind of overfitting via feature selection is a key motivation behind our use of the nested cross-validation approach (**Figure 2**): since the features are selected on the training and validation (test) set in the inner loop, any overfitting due to feature selection should not be transferred to the test set in the outer loop.

We have shown that only a few selected features seem to play a crucial role in classification throughout the cross-validation iterations (**Figure 5**).

We take a closer look at the 50 top features (which represent 4.24% of the total) comprised of 33 empirical features (17 Aβ, 12 Tau, four Volume) and 17 simulated features (**Supplementary Table D.2**). The anatomical distribution of feature relevance is summarized in **Figure 6**, for each, Aβ, Tau, and LFP.

**Figure 6.**
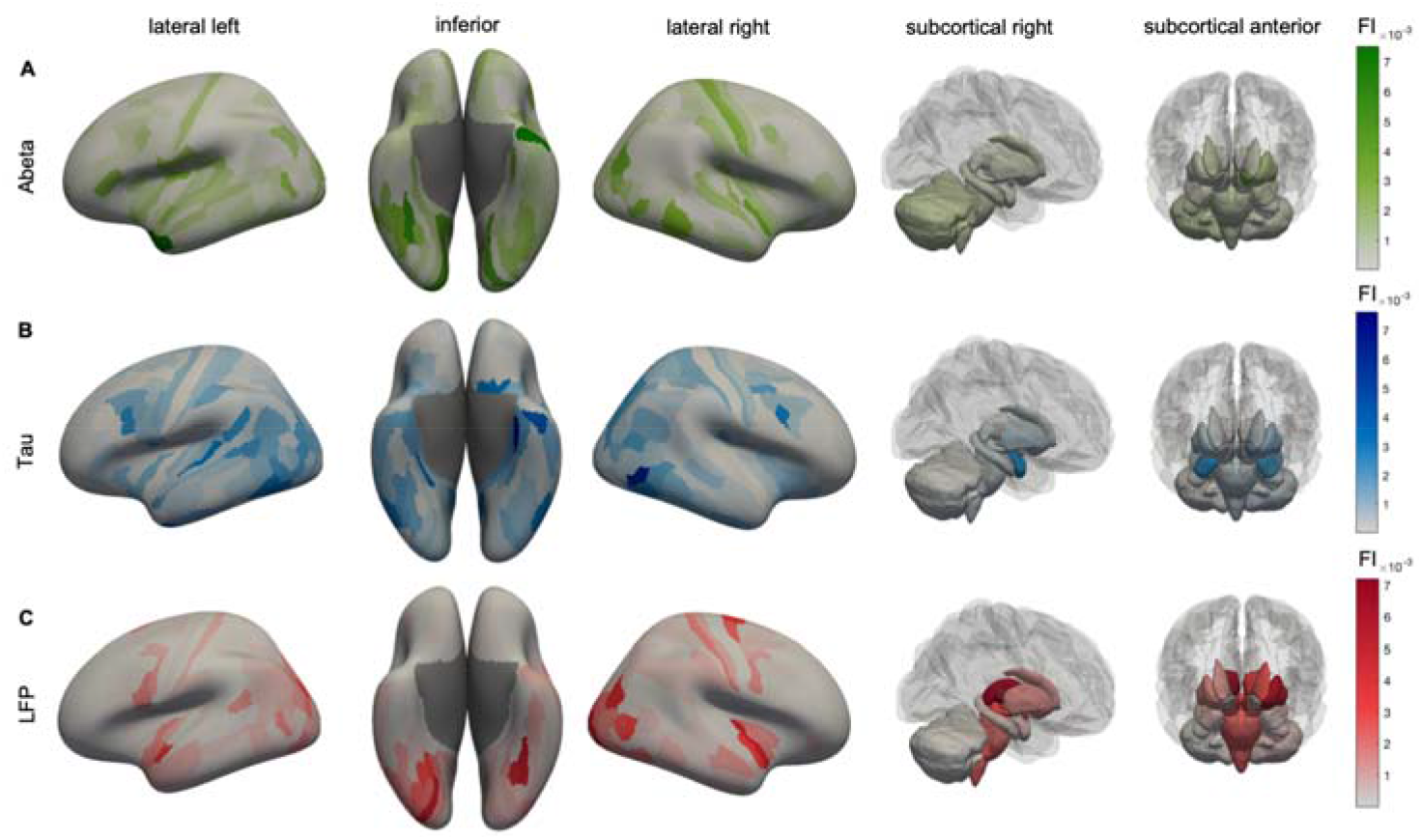
Anatomical representation of feature importance (FI) distribution. Displayed are cortical regions from left, right, and inferior as well as subcortical regions. The color indicates the feature importance. **(A)** Aβ FI. The anatomical patterns reveal high importance of left-temporal regions, as well as the left dorsal stream in the parietal and occipital cortex. The Aβ Top features showed a more disseminated allocation mostly in the temporal, occipital, frontal, and insular cortices, which is also in line with typical amyloid deposition and locations of increased AV-45 uptake in AD [37, 38]. (**B**) Tau features show a similar pattern as Aβ, but with a higher focus on typical Braak-stage-1 regions (as the entorhinal cortex). Most of the tau top features can be allocated to the temporal lobule, which is also the location of early Tau deposition according to the neuropathological BRAAK and Braak stages I-III [39-41] and the location of increased in-vivo binding of ^18^F-AV-1451 in AD [42, 43]. In particular, the entorhinal cortex is a consistent starting point of the sequential spread of Tau through the brain [39, 43] and also showed the most robust relationship between flortaucipir and memory scores in a recent machine learning study [44]. (**C**) Simulated frequencies do not show strong laterality as the empirical features but seem to have a focus in both occipital lobes, where typically alpha oscillations occur. The occipito-temporal and occipito-parietal regions of the first area are typical alpha-rhythm generators in resting-state EEG [45]. Alteration of these posterior alpha sources is a typical phenomenon in AD and MCI compared to HC [46]. The ventral or ‘what’ stream and the dorsal or ‘where’ stream have been implicated in object recognition and spatial localization and are essential for accurate visuospatial navigation [47, 48]. Impairment in visuospatial navigation is a potential cognitive marker in early AD/MCI that could be more specific than episodic memory or attention deficits [49, 50]. Besides this, subcortical areas like the thalami play a more crucial role than for Aβ and Tau.

The interpretation of the simulated features is more complicated and needs to be treated carefully.

In general, the simulated features depend on network information. Concurrently, in this work, all subjects used the same SC. Consequently, the difference between subjects can be attributed to the spatial distribution of their respective Aβ PET.

Many of the top simulated features can be allocated to two functional systems. Firstly, the visual cortex including the ventral and dorsal stream, and, secondly, subcortical structures like the thalamus and the putamen.

As a limitation of our study, we see that the used simulated feature, the mean simulated LFP frequency (averaged across a wide range of the large-scale coupling parameter G), is not directly equivalent to a biophysical measurement like empirically measured LFP. LFP frequency was averaged across the global scaling factor G. G scales the strength of long-range connections in the brain network model and is a crucial factor in the simulation. Many different dynamics can develop across the dimension of G, from which some are similar to empirically observed phenomena, but others are not. Our former work has found that particular ranges of G with non-plausible frequency patterns hold the potential to differentiate between diagnosis groups [17]. This is mainly because of the underlying mathematics of the Jansen-Rit model: besides two limit cycles that produce alpha-like and theta-like activity, the local dynamic model has a region of stable focus wherein no oscillations are produced in the absence of noise. Technically, this stable focus is represented as a zero-line artifact that appears mainly in the HC group, because only Aβ values above a critical value led to the presence of the slower theta-limit-cycle. By averaging LFP frequencies across the whole spectrum of G, we incorporate this zero-line information, which leads to apparently higher mean LFP frequencies for the AD group compared to non-AD groups. In contrast, in the region of biologically plausible results, AD has lower frequencies – as would be expected [17]. This can also be seen as another advantage of TVB. It shows how TVB does not just reproduce data that could also be obtained with EEG or intracranial electrodes, but delivers ‘artificial’ data that is still informative. While particular parameter ranges deliver biologically plausible results, even other (less plausible) parameter settings provide unique individual patterns and can contribute to the classification.

This work’s primary aim is not to develop a ready-to-use ML classifier for AD, but to show the potential of brain simulation to enhance empirical datasets in clinically relevant ways. While the limited sample size used in this study would potentially be problematic in a more traditional ML study aimed at providing an ML-based diagnostic aid, combined with our careful cross-validation methodology, it does not detract from our primary conclusion. Future studies will have to reproduce these results using a more extensive cohort for further clinical usage of this work. Ideally, external validation with a dataset outside of ADNI would be performed.

We used ML as an approach for the comparison of classifier performance with empirical data against simulated data, which is wholly derived from the empirical data. Improvement in classification is then strong evidence for successful processing of the empirical data in TVB – TVB decodes the information embedded within the empirical data, that cannot be detected by statistics or ML classifiers. We showed in ADNI data that TVB can derive additional information out of the spatial distribution pattern in PET images.

Our work provides novel evidence that TVB can act as a biophysical brain model - and not just like a *black box*. Complex multi-scale brain simulation in TVB can lead to additional information, that goes beyond the implemented empirical data. Our analysis of feature importance supports this hypothesis, as the features with the highest relevance are already well-known AD factors and hence, biologically plausible surrogates for clinically relevant information in the data. Moreover, in this pilot study, we demonstrate that TVB simulation can lead to an improved diagnostic value of empirical data and might become a clinically relevant tool.

## ^†^ ABBREVIATIONS

AD: Alzheimer’s Disease
Aβ: Amyloid β
TVB: The Virtual Brain
EEG: electroencephalography
LFPs: local field potentials
PET: positron emission tomography
MCI: mild cognitive impairment
HC: healthy controls
ML: machine learning
ADNI: Alzheimer’s Disease Neuroimaging Initiative
MRI: magnetic resonance imaging
SC: structural connectivity
SVM: support vector machines
RF: random forest
SUVR: standardized uptake value ratio
FI: feature importance

## APPENDICES

A. Image Processing
B. Electrophysiological Simulation with The Virtual Brain
C. Machine Learning Methodology
D. Supplementary Results

## AUTHOR CONTRIBUTIONS

All authors have made substantial intellectual contributions to this work and approved it for publication. PT and LS had equal contributions to this work. Particular roles according CrediT [36]:

**Paul Triebkorn**: Conceptualization, Data curation, Investigation, Methodology, Visualization, Writing - original draft. **Leon Stefanovski**: Conceptualization, Formal analysis, Investigation, Methodology, Visualization, Writing - original draft. **Kiret Dhindsa**: Formal analysis, Methodology, Software, Writing - review & editing. **Margarita-Arimatea Diaz-Cortes**: Methodology, Software, Writing - review & editing. **Patrik Bey**: Methodology, Software, Writing - review & editing. **Konstantin Bülau**: Validation, Writing - review & editing. **Roopa Pai**: Data curation, Writing - review & editing. **Andreas Spiegler**: Methodology, Writing - review & editing. **Ana Solodkin**: Writing - review & editing. **Viktor Jirsa**: Writing - review & editing. **Anthony Randal McIntosh**: Writing - review & editing. **Petra Ritter**: Conceptualization, Funding acquisition, Methodology, Project administration, Supervision, Writing - review & editing.

## DATA AVAILABILITY

The raw data for this study is available in ADNI. The codes used in this study are available on request to the corresponding author.

## ETHICS STATEMENT

The study providing the data for this work has been approved from the Ethics Board of the Charité - Universita□tsmedizin Berlin under the approval number EA2/100/19.

The work described in this article has been carried out in accordance with The Code of Ethics of the World Medical Association (Declaration of Helsinki) for experiments involving humans:

(http://www.wma.net/en/30publications/10policies/b3/index.html); the EC Directive 86/609/EEC for animal experiments:

(http://ec.europa.eu/environment/chemicals/lab_animals/legislation_en.html); and the Uniform Requirements for manuscripts submitted to Biomedical journals (http://www.icmje.org).

## ACKNOWLEDGMENTS

Data collection and sharing for this project was funded by the Alzheimer’s Disease Neuroimaging Initiative (ADNI) (National Institutes of Health Grant U01 AG024904) and DOD ADNI (Department of Defense award number W81XWH-12-2-0012). ADNI is funded by the National Institute on Aging, the National Institute of Biomedical Imaging and Bioengineering, and through generous contributions from the following: AbbVie, Alzheimer’s Association; Alzheimer’s Drug Discovery Foundation; Araclon Biotech; BioClinica, Inc.; Biogen; Bristol-Myers Squibb Company; CereSpir, Inc.; Cogstate; Eisai Inc.; Elan Pharmaceuticals, Inc.; Eli Lilly and Company; EuroImmun; F. Hoffmann-La Roche Ltd and its affiliated company Genentech, Inc.; Fujirebio; GE Healthcare; IXICO Ltd.; Janssen Alzheimer Immunotherapy Research & Development, LLC.; Johnson & Johnson Pharmaceutical Research & Development LLC.; Lumosity; Lundbeck; Merck & Co., Inc.; Meso Scale Diagnostics, LLC.; NeuroRx Research; Neurotrack Technologies; Novartis Pharmaceuticals Corporation; Pfizer Inc.; Piramal Imaging; Servier; Takeda Pharmaceutical Company; and Transition Therapeutics. The Canadian Institutes of Health Research is providing funds to support ADNI clinical sites in Canada. Private sector contributions are facilitated by the Foundation for the National Institutes of Health (www.fnih.org). The grantee organization is the Northern California Institute for Research and Education, and the study is coordinated by the Alzheimer’s Therapeutic Research Institute at the University of Southern California. ADNI data are disseminated by the Laboratory for Neuro Imaging at the University of Southern California.

Computation of underlying data has been performed on the HPC for Research cluster of the Berlin Institute of Health.

