## Appendix A for "Brain simulation augments machine-learning-based classification of dementia"

**Appendix A – Image processing**

**A.1. Data description**

We used data from ADNI in recent work to simulate electrophysiological neuronal activity (EEG) and explore their spectral characteristics in AD, MCI, and HC [1]. In the present study, we assess whether the results of the previous study can be used to improve ML classification of AD, MCI, and HC. Basic epidemiological properties can be found in [1] and are also presented in **Table A.1**.

**Table A.1.** Basic epidemiological properties of the included participants. Data taken from [1] with permission.

| Diagnosis | *n* (female) | Mean age | *σ* | Min. age | Max. age | Mean MMSE | *σ* | Min. MMSE | Max. MMSE |
| --- | --- | --- | --- | --- | --- | --- | --- | --- | --- |
| AD | 10 (5) | 72.0 | 9.6 | 55.9 | 86.1 | 21.3 | 6.8 | 9 | 30 |
| HC | 15 (9) | 70.6 | 4.7 | 63.1 | 78.0 | 29.3 | 0.8 | 28 | 30 |
| MCI | 8 (3) | 68.2 | 6.4 | 57.8 | 76.6 | 27.1 | 1.6 | 25 | 30 |

**A.2. Image processing**

In the following, we will provide a summary of data processing. We used the human connectome project (HCP) minimal preprocessing pipeline [2] for the processing of structural data. This included Freesurfer [3], FSL [4-6] and connectome workbench 7].

The adjustments on the HCP guidelines to fit our data are described in the methods of [1].

For detailed metadata descriptions, please compare Supplementary Table S1-S5 in [1]. We included T1 MPRAGE (TE = 2.95 - 2.98 ms, TR = 2.3s), FLAIR (TE differs slightly, TR = 4.8s, matrix size = 160 x 256 x 256), DWI (TE = 56 -71 ms, TR = 3.4 - 7.2s, matrix size = 116 x 116 x 80, voxel size = 2 x 2 x 2, bvals = [0, 1000] or [0, 500, 1000, 2000], bvecs = 49 or 115), fieldmaps and PET Data (AV-45 for Aβ and AV-1451 for Tau).

For AV-45 and AV-1451 PET images, we used the already preprocessed images available in ADNI. We aligned the PET images to HCP-processed T1 images and performed linear registration with FLIRT (FSL). The resulting PET images were masked with subject-specific HCP brain masks. We calculated SUVRs by dividing image intensities by the mean intensity in the white matter of the cerebellum.  Partial volume correction was applied using grey and white matter from Freesurfer segmentation and the Müller-Gärtner method from the PETPVC toolbox [8].   To get the average SUVR per region, subcortical SUVRs were defined as the average SUVR in subcortical GM. Simultaneously, cortical GM PET intensities were mapped onto the cortical surfaces using the connectome workbench tool. DWI preprocessing was performed by the MRtrix3 software package (<http://www.mrtrix.org>). We used the following functions: *Dwidenoise* [9]*, Dwipreproc* [10]*, Dwibiascorrect*, *Diw2mask, Dwiintensitynorm, Dwi2response* [11]*, Average_response, Dwi2fod*, [12], *Tckgen* [13-15]. The detailed steps are described in the original paper [1]. The preprocessed cortical surfaces and T1 images were used to compute the Boundary Element Model in Brainstorm [16], wherein inner and outer Scalp, as well as outer skull, were modeled with 1922 vertices per layer and the default 'BrainProducts EasyCap 65' EEG cap. We estimated the leadfield matrix with the adjoint method in OpenMEEG (default conductivities 1 (scalp), 0.0125 (skull), and 1 (brain)).
