## Appendix B for "Brain simulation augments machine-learning-based classification of dementia"

**Appendix B - Electrophysiological Simulation with The Virtual Brain**

As simulated features for ML classification, we use simulation results of [1] originally done with Aβ PET SUVR from ADNI. A detailed description of the corresponding equations can be found there.

In TVB, the long-range input to a region *i*, induced by the output 𝑥_𝑗_ from a region 𝑗, developes along the structural connectivity matrix 𝑆𝐶_𝑖𝑗_ (connectivity between regions *i* and *j).* It is scaled by the global scaling factor *G* and generally defined as:

Eq. B.1

$\dot{x_{i}}=N\left( x_{i}\left( t \right) \right)+G\sum_{1}^{n} SC_{ij}x_{j}$

Wherein $N\left( x_{i}\left( t \right) \right)$the local dynamic model function and $\dot{x}_{i}$ the derivative of its state variable. We explored a range of 0 < G < 600 in steps of 3, leading to 201 LFP values per region. This allows to explore different dynamic behaviors, in particular as former studies have identified the global scaling parameter G as a crucial factor for biologically plausible results [2, 3]. The LFP was averaged over G for each of the 379 regions to be used as simulated features for ML classification. This average is able to capture different “dynamic states” of the brain in one value per region [2].

As the local dynamic model, we employ the Jansen-Rit neural mass model [4-6], considering the three neural populations of pyramidal cells, inhibitory interneurons, and excitatory interneurons. The Jansen-Rit model has been shown particular abilities to reproduce biologically plausible frequency spectra in electroencephalography [4]. The membrane potential and electrical behavior for the three cell types is described by two state variables for each population, namely y_0_ – y_5._ They are defined in six differential equations together with their derivatives $\dot{\boldsymbol{y}}$_0_ - $\dot{\boldsymbol{y}}$_5_:

Eqs. B.2 – B.7

$$\dot{y_{0}}=y_{3}$$

$$\dot{y_{3}}=Aa S\left[ y_{1}-y_{2} \right]-2 ay_{3}-a^{2} y_{0}$$

$$\dot{y_{1}}=y_{4}$$

$$\dot{y_{4}}=Aa \left[ p\left( t \right)+\alpha_{2}JS\left[ \alpha_{1}J y_{0} \right]+c_{0} \right]-2ay_{4}-a^{2} y_{1}$$

$$\dot{y_{2}}=y_{5}$$

$$\dot{y_{5}}=Bb\left( \alpha_{4}J S\left[ \alpha_{3}J y_{0} \right] \right)-2by_{5} -b^{2} y_{2}$$

Wherein *A* and *B* are excitatory and inhibitory amplitudes, and *a* and *b* are the corresponding dendritic time constants; *J*, $\alpha$*_1_*, $\alpha$*_2_*, $\alpha$*_3_*, and $\alpha$*_4_* are coupling parameters between the neural populations; *p(t)* is the input noise and *c_0_* is the input from connected regions.

The membrane potential $v$ is translated into a firing rate $S\left[ v \right]$ by the potential-to-rate function:

Eq. B.8

$$S\left[ v \right]=\frac{2 \nu_{max}}{1+e^{r\left( v_{0}-v \right)}}$$

Wherein the maximum firing rate is 2𝜈_𝑚𝑎𝑥_, the steepness parameter *r*, and inflection point 𝑣_0_.

The LFPs are estimated as the difference $y_{0}- y_{2}$ between state variables of pyramidal cells and inhibitory interneurons. This reflects the outgoing electrical activity from (projecting) pyramidal cells, reduced by local inhibitory effects.

The complete implementation of the Jansen-Rit model in TVB is openly available. The source code can be found here:

<https://github.com/the-virtual-brain/tvb-root/blob/master/scientific_library/tvb/simulator/models/jansen_rit.py>

To locally adjust the inhibitory time constant *b* by Aβ concentration *x*, we introduced a sigmoidal transfer function *b(x)*:

Eq. B.9

$$\left\{ \begin{aligned} x_{0}=\frac{(x_{2}-x_{1})}{2}+x_{1} \\ k=\frac{\ln(\frac{y_{max}}{c}-1)}{x_{2}-x_{0}} \\ b\left( x \right)=\frac{y_{max}}{1+e^{k\cdot(x-x_{0})}}+y_{min} \end{aligned} \right.$$

Wherein y_min_ is the lower asymptote of the sigmoid; y_min_ + y_max_ is the upper asymptote; x_1_ and x_2_ describe the range of Aβ SUVR wherein the time constant *b* changes; *k* and c adjust the curve steepness; and x_0_ is the inflection point of the sigmoid.
