## Appendix C for "Brain simulation augments machine-learning-based classification of dementia"

**Appendix C – Machine Learning Methodology**

**C.1. Background**

Over the last 15 years, ML has evolved as a fast-growing multidisciplinary field of research, with many applications in various scientific areas [1], including neuroimaging in AD [2]. ML techniques range from relatively simple mathematical models to complex approaches. Their principal aim has been to “give computers the ability to learn without being explicitly programmed to do so” [3]. In other words, ML can be seen as a collection of methods developed to enable computational systems to learn from the data with the primary purpose of making predictions and inferences. The advantage of ML algorithms over traditional statistical or model-based approaches is that they can discover subtle and even complex patterns in high-dimensional data that would be difficult to identify or encode otherwise [4]. However, the ability of ML algorithms to discover patterns in data can also result in the reliance on spurious correlations that appear in data by chance, or which are otherwise not clinically generalizable beyond the data used to train the algorithm. Many algorithms have been developed with successful convincing classification capabilities [5, 6].

Generally, the problem of classification is the prediction of categories for each point in a (multidimensional) data cloud based on their representations, that is, identifying the class to which an input belongs among a set of labeled categories [7]. A classification model, also called a classifier, can be binary or multi-class depending on the number of groups or labels to predict. A classifier is then tailored from the learning process of categorizing a set of “training” data. Despite the numerous classification methods currently available for this inference, it is impossible to conclude which one is generally superior to the other. This particular fact depends on the application and nature of the available data set and is commonly referred to as the “no-free-lunch theorem" [8].

Classification allows considering other factors, like lifestyle risks, genetic conditions, among others [9]. Numerous ML methods have been used to classify and predict AD stages with promising results [10-12]. While some studies have made use of a single screening modality, such as MRI [13-17], or electroencephalography (EEG) [18-21], others have used a combination of multiple imaging techniques including MRI, PET, and cerebrospinal fluid (CSF) biomarkers [22-28]. Although many of those studies presented interesting and promising results in AD classification, most focused on a so-called two-class problem. It has been pointed out, that standardization and quality checks for robustness, CV schema, etc. are essential but often lacking [21]. Only a few studies have made use of more complex classification, as a three-class problem with AD, MCI, and HC [20, 29] or differentiation in between MCI to converters and non-converters [30-33].

**C.2. Support Vector Machines**

Among the spectrum of ML classifiers, in the present study, we developed a dual methodology for solving the classification task, by using Support Vector Machine (SVM) and Random Forest (RF). Both SVM [12, 34-37] and RF (for a review see [38]) are established ML classifiers in neuroscience. Both approaches were cross-validated within a sample of 33 human subjects with MCI, AD, or HC from the ADNI database. The use of two different ML-classifiers in a nested approach provides a robust, generalizable, and appropriate evaluation of the classification; on the other hand, it enables exploring the empirical and simulated features of highest importance for separation between the three groups under study.

SVM is a widely used method for supervised ML classification problems and has been well established in recent neuroscience literature [12, 34-37]. SVMs have been extensively employed due to their robustness, simplicity to implement, and because they can also be employed as non-linear classifiers by making simple variations. SVM is a specific type of so-called maximum margin classifiers that tries to find an optimal separating hyperplane (with the largest possible margin) between two (or more) groups within a higher dimensional representation of the original data. An SVM aims to define a decision-boundary, or a set of boundaries in the case of multi-class classification, which partitions the feature space into class-defining regions. In the very simple case of only two features and two classes, this would mean separating two clusters of points on a two-dimensional plot with a line. This is done by maximizing the margins between boundaries, which are referred to as the support vectors. When this is not possible in the original feature space, non-linearity is introduced via the SVM’s kernel function. This function is used to project the data into an arbitrarily high dimensional space wherein the best decision boundary is linear (i.e., a hyperplane). The projection is then reversed so that the decision boundary can be projected back into the original feature space, resulting in a non-linear boundary.

Due to its algorithm particularities, SVM has some advantages in small sample analysis [39], which in this approach represents an advantage since we employ a sample size of 33 subjects.

In the following, we explain the hyperparameters (i.e., the parameters that describe characteristics of how the classifier works) that were adjusted for SVM in this study.

**C.2.1. Kernel function**

The kernel of an SVM is the function that is used for the transformation of the feature space. A kernel is an invertible function $k\left( x_{i},x_{j} \right)=r\left( x_{i} \right)\cdot r\left( x_{j} \right)$ used to transform data points $x_{i},i\in1\ldots N$ in such a way as to preserve the relationships between datapoints so that the SVM can learn a linear classification rule in the feature space of $r\left( x_{i} \right)$ that corresponds to a non-linear classification rule in the feature space of $x_{i}$. The classification rule is learned in the transformed space and then projected back down into the original data space to obtain a non-linear classification boundary. I.e., the kernel is the function that transforms the original feature space into a higher dimensional (artificial) feature space to separate data points, while it also defines the rule for its back-projection.

We explored two types of kernels. The Gaussian radial basis function (RBF) kernel is the most commonly used kernel for non-linear problems for a variety of reasons (e.g., the kernel is stationary, smooth, and tunable with only a single isotropic parameter). The kernel is defined as follows: $k\left( x_{i},x_{j} \right)=exp\left( -\gamma\left| \left| x_{i}-x_{j} \right| \right|^{2} \right)$ for $\gamma>0.$ The polynomial kernel instead raises the degree of the data space using, i.e., quadratic or cubic transforms: $k\left( x_{i},x_{j} \right)=\left( x_{i},x_{j} \right)^{d}$. Here we only explored $d\in2,3,4$ to avoid overly complex decision functions.

**C.2.2. Scaling parameter Gamma**

The scaling parameter as defined in the RBF kernel (see above). It scales the distance of datapoints to the decision boundary that are used for its calculation. This must typically be tuned empirically using cross-validation, as it can take any value over zero, including values less than one to shrink the norms between points, or values much larger than 1, to expand those norms, depending on how the data are clustered.

**C.2.3. Soft-margin parameter C**

The soft-margin parameter C controls whether the solution emphasizes a wide margin (i.e., a decision boundary as far away as possible from the closest points), which can lead to some underfitting, versus a narrow margin that can result in some overfitting. E.g., an extremely narrow margin around zero could still perfectly separate datapoints of the training set, but as the decision boundary almost crosses the most similar datapoints between the classes, it will not be very adaptable to new data. Like Gamma, it must be tuned empirically.

**C.2.4. Number of Features**

Since our list of candidate features is very high compared to the number of subjects, to find a more robust and interpretable solution [40, 41] , we greatly limit the number of selected features prior to classification.

For the SVM parameters Gamma and C, these are fairly standard search ranges [42, 43]. Typically a search will span different orders of magnitude, for example in base 10 or in base 2. We used a coarse grid in our hyperparameter search so as to not overdetermine our model to our relatively small sample.

**C.3. Random Forests**

Similar to SVMs, Random Forest (RF) has been widely used for classification within the ML community and has performed well in a range of applications for classification in neuroimaging studies [38]. It is based on a large number of decision trees performing binary splits on randomly selected subsets of features, and therefore uses a fundamentally different classification technique than SVMs. An RF builds many decision trees based on finding the best partitioning of random subsets of features. Each one of those decision trees is constructed by a set of rules, learned from the data, organized in a hierarchy that determines the decision process for classification. So-called hyperparameters of the RF define how the trees should be constructed, e.g., how many layers they have or how many features are involved in each layer. Each data point is classified by each tree based on the path it travels, given the values of each feature. Each tree is then given a ‘vote’ on how to classify the datapoint. Finally, a classification decision of the RF is made by pooling all of the trees' decisions.

The main advantages are the resulting direct interpretability of the feature importance and high robustness towards overfitting of the algorithm [44]. One of the main differences between RF and SVM is that RF directly provides a probability for each point of belonging to a defined class. In contrast, SVM provides the distance to the hyperplanes or boundaries.

In the following, we explain the hyperparameters that were adjusted for the RF.

**C.3.1. Class Weight**

Simply whether or not the model should take into account the imbalanced class representation in the training set. When balanced, incorrect class labels during the training process are penalized more or less depending on whether the correct class is under- or over-represented in the training set. This aims to overcome biases that arrive from different frequencies of classes in the underlying training data.

**C.3.2. Number of Estimators**

The number of decision trees to train, i.e. the class estimates of which are aggregated in the final model. In other words, the size of the ensemble.

**C.3.3. Minimum samples per split**

The minimum number of divergent samples required to split a branch of a tree into two new branches. A lower value means a more detailed tree that can be more precise. E.g., the lowest value would be 1, meaning that even a single subject can be separated by a decision rule of one tree. Lower values are typically required when the number of samples are low, especially compared to the number of features.

**C.3.4. Minimum samples per leaf**

The minimum number of samples required at the end of each leaf node (the last node of each path down the tree). Similar to min. samples per split, this influences how detailed the tree can become, and lower values are typically required for a low sample setting with many features. When a branch reaches this number, it can no longer be split further.

**C.3.5. Maximum features**

The maximum number of features to consider when evaluating the best split for each branch. Lower values typically mean better generalization but reduced flexibility.

For the RF parameters, we kept to low values for splitting criteria so that more detailed trees could be learned (as mentioned, this is more important for our N<<P scenario, since generalization is much harder).

**C.4. Feature Space**

Prior to using the features in the machine learning procedure, we checked them manually for general plausibility. This led us to remove five Volume features from the freesurfer volumetrics outcome: ventricle and white matter hyperintensity measurements (5th Ventricle, left WM hyperintensities, right WM hyperintensities, left non-WM hyperintensities, and right non-WM hyperintensities). For technical reasons, these volumes were not obtained in the image preprocessing.

In the end, we used the following feature spaces:

1. Empirical features (800 dimensions)
   a. 379 values for regional Aβ burden in SUVR, measured from AV-45 PET
   b. One value for the averaged global Aβ burden
   c. 379 values for regional Tau burden in SUVR, measured from AV-1451 PET
   d. One value for the averaged global Tau burden
   e. 40 Volume measures from HCP standards image processing
2. Simulated features (379 dimensions)
   a. 379 regional LFP peak frequencies, averaged over 201 simulations with different scaling factor G
3. Combined features (1179 dimensions)
   a. All features from above

**C.5. Nested Cross-Validation Scheme**

We used a part of the population with subjects of all groups as a training population while providing specific data aspects to the machine-learning engine. For an overview of machine learning approaches on AD MRI and PET, see [24]. Our approach uses a more complex cross-validation with an inner and an outer cross-validation loop.

Our nested cross-validation scheme is illustrated in **Figure 2** of the main manuscript. By nesting two cross-validation loops, we are able to simultaneously perform feature selection and model selection robustly. Subjects are portioned into training and test sets in the main outer loop. The training set is then treated as if it were the full dataset in the inner cross-validation loop, where it is splitted again into a smaller training set and a validation set. In this inner loop, each parameter setting of the classifier (**Supplementary Tables C.1 and C.2**) undergoes 10-fold cross-validation with feature selection performed independently on each run. These variations on the model can then be compared by their performance against the validation set. The best performing model parameters are then used to train a new classifier on the larger training set defined in the outer loop, with performance measured on the actual test set. The features selected in the outer loop are stored to measure the frequency of selection for each feature. This entire process is repeated 100 times in order to obtain statistically reliable estimates of our chosen performance metrics and feature importance metrics. This is also why we use random sampling with replacement to partition training and test data since it allows a greater number of cross-validation iterations for statistical evaluation of our models.

The inner cross-validation loop is the model selection loop wherein hyperparameters can be optimized without influence from the test set to ensure an unbiased estimate of model generalization performance. The outer cross-validation loop is the standard cross-validation loop used to estimate model generalization performance using training and test set partitioning. Using a nested cross-validation loop to separate hyperparameter optimization and a model performance optimization is a well-documented approach to ensuring the validity of machine learning results and preventing overfitting [45, 46]. This is particularly necessary for the problem presented in this work, where the number of subjects is low, and the number of features is relatively high. Under such conditions, both overfitting and underfitting due to a poor choice in hyperparameters can significantly affect model performance. Therefore more robust methods like nested cross-validation are required.

Since N<<P and the complexity of the problem are such that we expect the possibility of considerable individual differences among the patient population, the influence of a single data point can be significant. This means learning a classification rule that is nearly optimal for one problem is quite difficult unless hyperparameter tuning is done. Technically, hyperparameter tuning should always be done for a "final model". Still, for more straightforward problems with good data representation, the influence of hyperparameters should not be as large as they are in our situation. So hyperparameter tuning is more a necessity due to the complexity of our problem. The validity of our results comes from the nested cross-validation loop we used, the way we search hyperparameters to prefer less complex and therefore more general solutions (e.g., using fewer features), and the empirical plausibility of the features that were selected. In other words, we performed the necessary step of hyperparameter optimization cautiously and conservatively to ensure the validity of our results.

**Supplementary Table C.1.**

Best hyperparameter settings for each feature set for SVM only.

| **Parameter** | **Empirical** | **Simulated** | **Combined** | **Searched parameter space** |
| --- | --- | --- | --- | --- |
| Kernel | RBF | RBF | Polynomial (d=3) | Radial basis function (RBF), polynomial functions |
| Gamma | 0.01 | 0.1 | n.a. | Gamma ∈ {10^-3^, 10^-2^, 10^-1^, 1} |
| C | 1000 | 100 | 1000 | C ∈ {10^-2^, 10^-1^, 1, 10, 10^2^, 10^3^} |
| Number of features K | 30 | 10 | 40 | X ∈ {5, 10, 15, 20, 25, 30, 35, 40} |

**Supplementary Table C.2.**

Best hyperparameter settings for each feature set for RF only.

| **Parameter** | **Empirical** | **Simulated** | **Combined** | **Searched parameter space** |
| --- | --- | --- | --- | --- |
| Class weight | balanced | balanced | balanced | None or balanced |
| Number of Estimators | 10 | 10 | 10 | n ∈ {10, 50, 100, 200} |
| Min. samples per split | 2 | 2 | 4 | n ∈ {2, 3, 4, 5} |
| Min. samples per leaf | 1 | 3 | 2 | n ∈ {1, 2, 3} |
| Max. features | $\sqrt{P}$ | none | $\sqrt{P}$ | n ∈ {P^-1/2^, log_2_P} |

**C.6. Feature selection methods**

For each classifier, we used an appropriate feature selection method:

To select features for the SVM classifier, we ranked features in the training set according to their F-statistic, as in an ANOVA analysis, which estimates the linear dependence of each feature on the class labels. We chose the top k features for varying values of k as part of the model selection phase of our nested cross-validation loop (**Supplementary** **Table B.1** for values).

RFs incorporate their own embedded feature selection process. Here we used entropy to allow the classifier to rank features by information gain while building trees. The maximum number of allowable features was the square root of the total number of input features, 34.

The entropy criterion uses the notion of entropy from Shannon’s Information Theory [47] as a measure of feature importance, i.e., by measuring how well it separates classes.

Eq. C.1

$$\text{Entropy}=\sum_{i=1}^{C} {-f}_{i}\text{log}\left( f_{i} \right)$$

Where $f_{i}$ is the proportion of class label $i$ that meets a splitting criterion learned for a given feature (e.g., the proportion of class label $i$ for which the feature is less than a particular value), and C is the number of classes.

**C.7.** **Classification Experiments**

Since our overall goal is to assess whether the inclusion of features extracted from TVB simulations contributes diagnostic information independent from just the empirical features, we repeat the entire machine learning process with three feature sets: 1) with the empirical features only; 2) with the simulated features only; 3) with both the simulated and empirical features combined.

To summarize, the design of our methodology posits:

1. a three-class task for AD, MCI, and HC with ML-classification
2. a nested dual classifier approach with SVM and RF
3. various sources of biological information in a “hybrid” methodology: multimodal empirical imaging data as well as simulated brain dynamics

In order to establish the contribution of our feature sets separately from the contribution of our classification approach, we completed nine experiments organized in a 3x3 grid (three feature sets by three classification approaches). As described above, the three feature sets used are the empirical features, the simulated features, and the combination of empirical and simulated features via concatenation. For our classification approaches, we use the SVM and RF approaches already described, as well as a combined approach. In the combined method, we use the RF for feature selection. Its embedded feature selection approach is well-designed for taking into account the interactions among large numbers of features instead of the univariate method used with the SVM. The SVM is used for classification due to its power in low-sample settings once a robust feature set is preselected, owed in part to its maximum-margin objective.

**C.8.** **Classification Performance Evaluation**

We used the weighted F1-score to evaluate classifier performance. The F-statistic ranks each feature based on their ANOVA F-value to measure class separability and select the top k features [48].

This metric is particularly beneficial for classification problems with imbalanced classes, where classification accuracy alone can be misleading (consider a classifier that only predicts Healthy Control with probability 1: it would have a 45.5% classification accuracy, which can be misinterpreted as better than chance for a 3-class problem). The weighted F1-score is computed by taking each class-wise F1-score, which itself is an average of precision and recall for the given class, and averaging those F1-scores together weighted by the number of samples in each class:

Eq. C.2

$$weighted F1=\frac{1}{N}\sum_{k\in K} fr\left( C_{k} \right)F1_{C_{k}}$$

where N is the total number of data samples, K is the number of classes represented by Ck, fr(Ck) is the frequency of class k, and

Eq. C.3

$$F1=2\frac{precision \cdot recall}{precision+recall}$$

For comparison of weighted F1-scores between the groups (empirical data, simulated data, and combined data in the feature space) we used the Wilcoxon Signed-Rank test, as the Shapiro-Wilk test revealed p < 0.01 for the empirical and combined approach (non-normal distributed) and p = 0.07 for the simulated approach (normal distributed), leading to the usage of a non-parametric test. We assessed the significance by using data from 100 cross-validation runs, leading to 100 data points per group.

**C.9.** **Feature Importance Metrics**

We tracked two metrics for feature importance. The most direct feature importance metric is the feature importance statistic used for feature selection itself, i.e., the F-statistic for the SVM and the entropy measure for the RF. In conjunction, we also tracked the selection frequency, defined as the proportion of outer cross-validation iterations in which each feature was selected. We compare and contrast these two metrics and discuss their agreement and differences below.

**C.10** **Performance validation by permutation testing**

The confidence intervals of classification performance values across folds in cross-validation schemes are typically large in small-sample settings, and present an issue for the interpretability of classification results [49].

For each permutation run, the resulting accuracy metrics of the 100 cross-validation loops were averaged to ensure comparability to the distribution of F1 scores of the actual model. We then compared the distribution of accuracies between the null models of each of the three approaches (empirical, simulation, combined) using Wilcoxon’s rank sum test to test for significance in model performance and relation to the CV based prediction error.
