## Appendix D for "Brain simulation augments machine-learning-based classification of dementia"

**Appendix D – Supplementary Results.**

**D.1. Classification results for all experiments**

A detailed description of the schemes with RF only and SVM only can be found in **Supplementary Tables C.1 and C.2**. For a more detailed visualization of the results of SVM only and RF only classification, consider **Supplementary Figures D.1 and D.2.** The overall weighted F1-score performances are presented in **Table D.1.**

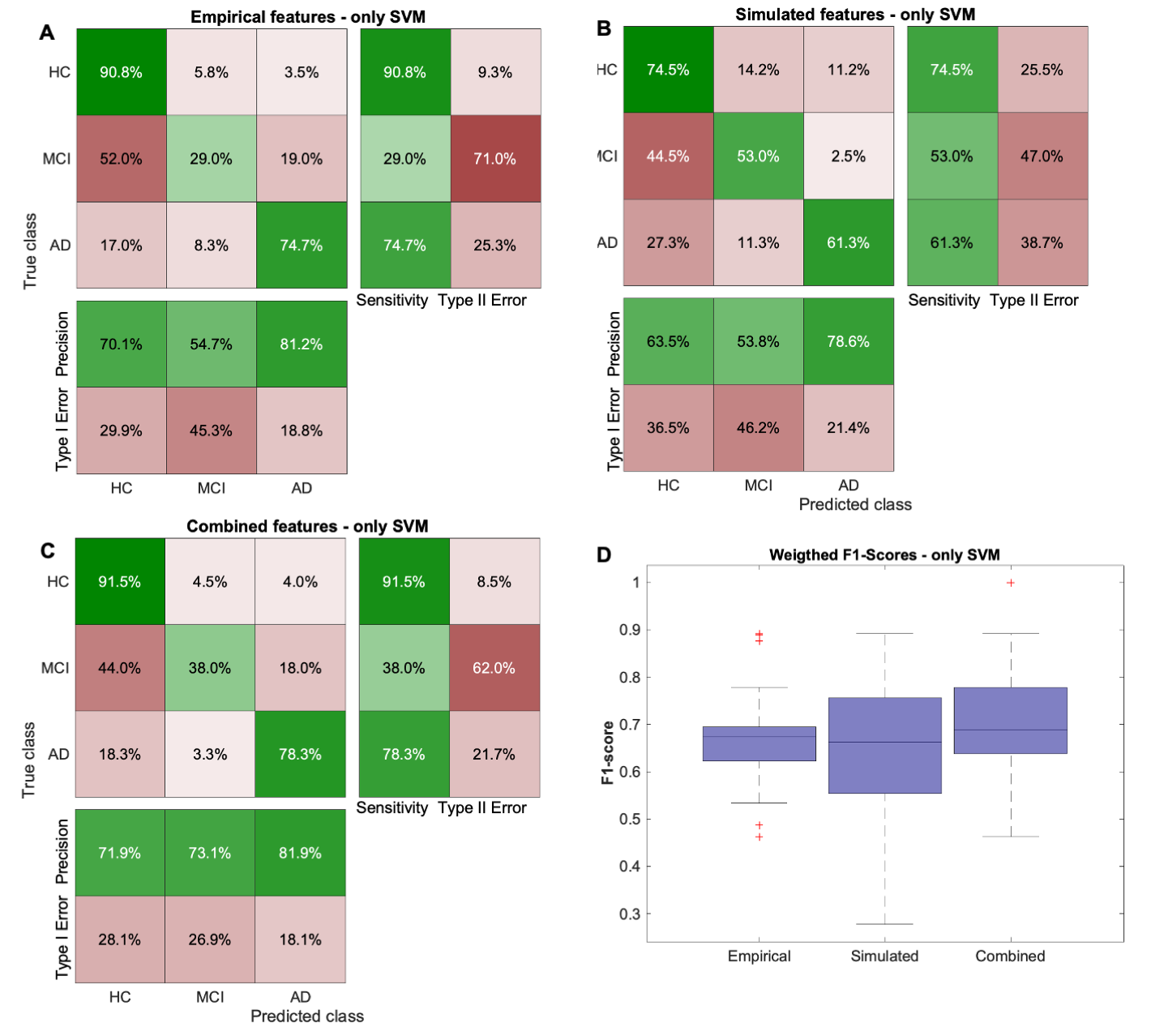

**Supplementary Figure D.1.** Results of SVM classification approach. (**A-C**) Confusion matrices are computed by summing the confusion matrices across all 100 cross-validation runs and normalizing per class. As in the (superior) nested approach mentioned in the main text, the combined approach improved the prediction of MCI participants. (**D**) Boxplots of mean F1-scores for three different feature spaces.

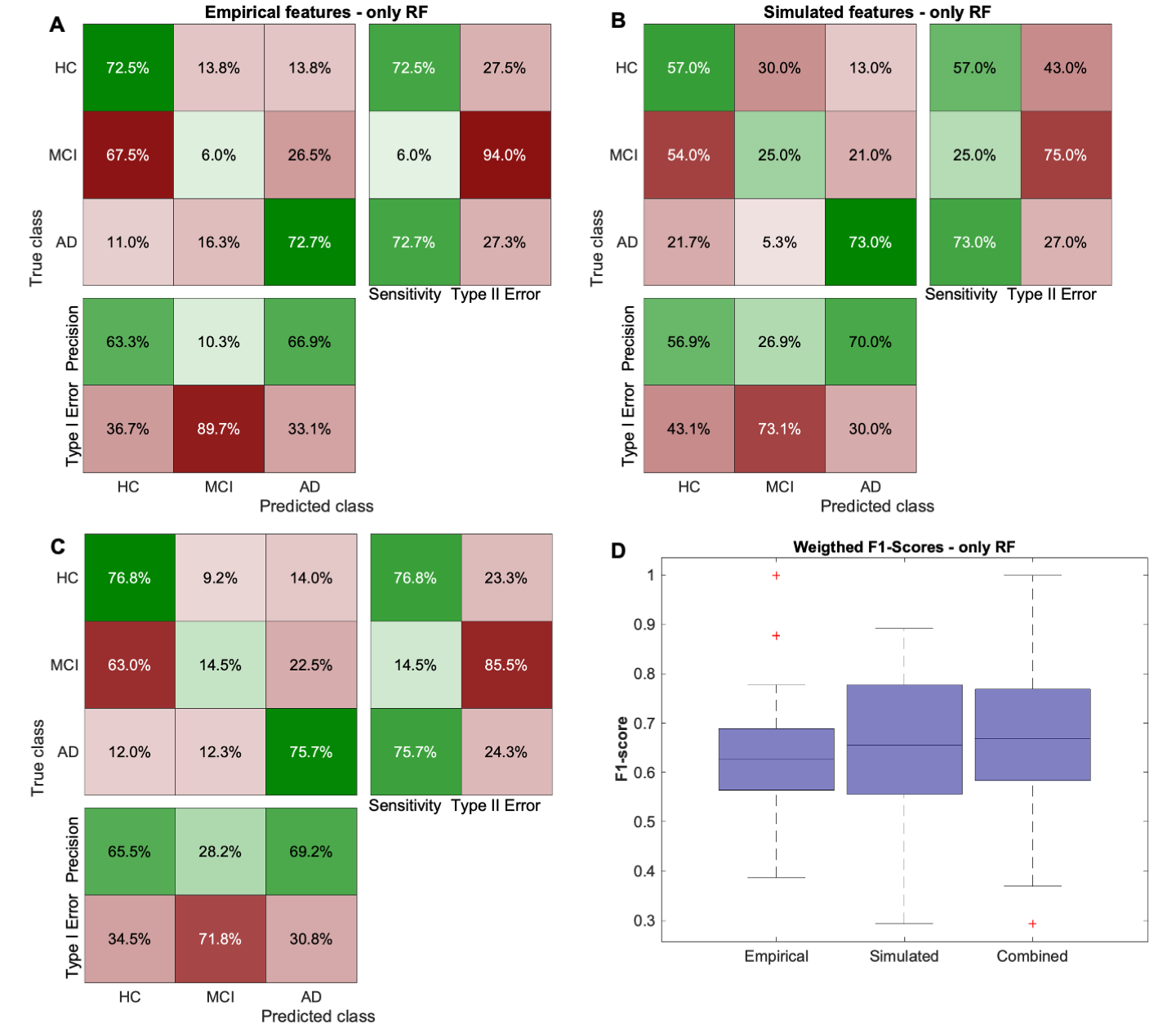

**Supplementary Figure D.2.** Results of RF classification approach. (**A-C**) Confusion matrices are computed by summing the confusion matrices across all 100 cross-validation runs and normalizing per class. (**D**) Boxplots of mean F1-scores for three different feature spaces.

**Table D.1.** Classification performance for different experimental designs.

| F1-score | SVM | RF | SVM + RF |
| --- | --- | --- | --- |
| Empirical features | 0.6756 | 0.6304 | 0.6434 |
| Simulated features | 0.6338 | 0.6501 | 0.6607 |
| Combined features | 0.7182 | 0.6699 | 0.7428 |

**D.2. Detailed analysis of feature importance**

**Table D.2** shows the 50 highest ranked features from all modalities.

**Table D.2.** **Full name description and functional network association of the 50 top features with the highest feature importance in the classification problem**. Parcellation adapted from [1]. Functional networks adapted from [2]. DS: Dorsal Stream, VS: Ventral Stream, SMA: Supplementary motor area

| **Rank** | **Feature Name** | **Full-Parcel-Name** | **Network** |
| --- | --- | --- | --- |
| 1 | L_EC_ROI_Tau: | Entorhinal cortex | Default mode |
| 2 | L_TGd_ROI_Abeta | Dorsal temporal gyrus | Default mode |
| 19 | R_PHA1_ROI_Tau | Parahippocampal area 1 | Default mode |
| 21 | L_POS1_ROI_Tau | Parieto-occipital sulcus area 1 | Default mode |
| 22 | L_EC_ROI_Abeta | Entorhinal cortex | Default mode |
| 26 | R_POS1_ROI_Abeta | Parieto-occipital sulcus area 1 | Default mode |
| 32 | L_pOFC_ROI_Tau | Posterior orbitofrontal cortex | Default mode |
| 4 | R_FST_ROI_Tau | Fundus of superior temporal sulcus | Visual |
| 8 | R_V3_ROI_LFP | Visual area 3 | Visual |
| 9 | L_V2_ROI_Abeta | Visual area 2 | Visual |
| 10 | R_V1_ROI_Abeta | Visual area 1 | Visual |
| 13 | R_ProS_ROI_Abeta | Prostriate region | Visual |
| 38 | R_FST_ROI_Abeta | Fundus of superior temporal sulcus | Visual |
| 39 | R_MST_ROI_LFP | Medial superior temporal area | Visual |
| 49 | L_MST_ROI_Abeta | Medial superior temporal area | Visual |
| 6 | R_VVC_ROI_Abeta | Ventral visual complex | Visual (VS) |
| 7 | L_VVC_ROI_LFP | Ventral visual complex | Visual (VS) |
| 11 | L_VMV3_ROI_LFP | Ventromedial visual complex 3 | Visual (VS) |
| 27 | R_FFC_ROI_Abeta | Fusiform face complex | Visual (VS) |
| 33 | R_VMV2_ROI_LFP | Ventromedial visual area 2 | Visual (VS) |
| 35 | L_PH_ROI_Tau | Area PH in lateral occipital lobe | Visual (VS) |
| 40 | R_PH_ROI_Tau | Area PH in lateral occipital lobe | Visual (VS) |
| 45 | R_VMV3_ROI_LFP | Ventromedial visual complex 3 | Visual (VS) |
| 47 | R_VMV3_ROI_Abeta | Ventromedial visual complex 3 | Visual (VS) |
| 42 | L_V3B_ROI_LFP | Visual area 3b | Visual (DS) |
| 16 | R_PGp_ROI_LFP | Parietal area G posterior | Dorsal attention |
| 17 | R_PGp_ROI_Abeta | Parietal area G posterior | Dorsal attention |
| 12 | R_PoI1_ROI_LFP | Posterior insula 1 | Cingulo-opercular |
| 18 | R_PI_ROI_Abeta | Parainsular cortex | Cingulo-opercular |
| 36 | R_PI_ROI_LFP | Parainsular cortex | Cingulo-opercular |
| 20 | L_TA2_ROI_LFP | Temporal region A | Auditory |
| 30 | L_PBelt_ROI_Tau | Parabelt complex (Auditory cortex) | Auditory |
| 28 | R_6mp_ROI_LFP | Area 6 medial posterior (SMA) | Somatomotor |
| 29 | L_Ig_ROI_Abeta | Insula granular cortex | Somatomotor |
| 34 | R_5m_ROI_Tau | Area 5 medial of paracentral lobule | Somatomotor |
| 44 | R_3b_ROI_Abeta | Area 3b of postcentral gyrus | Somatomotor |
| 14 | L_TGv_ROI_Tau | Ventral temporal gyrus | Language |
| 46 | L_45_ROI_Abeta | Area 45 of inferior frontal gyrus | Language |
| 31 | R_TE1p_ROI_Abeta | Temporal area 1 posterior | Frontoparietal |
| 37 | R_IFJp_ROI_Tau | Inferior frontal junction posterior | Frontoparietal |
| 3 | Right-Thalamus-Proper_LFP | Thalamus proper | Subcortical |
| 5 | Left-Putamen_LFP | Putamen | Subcortical |
| 23 | Left-Thalamus-Proper_LFP | Thalamus proper | Subcortical |
| 41 | Right-VentralDC_LFP | Ventral diencephalon | Subcortical |
| 43 | Right-Amygdala_Tau | Amygdala | Subcortical |
| 50 | Left-VentralDC_LFP | Ventral diencephalon | Subcortical |
| 15 | Right-Putamen_Volume | Putamen | Volume |
| 24 | Left-Hippocampus_Volume | Hippocampus volume | Volume |
| 25 | Right-Hippocampus_Volume | Hippocampus volume | Volume |
| 48 | Left-Caudate_Volume | Nucleus caudatus | Volume |

In the following, we further evaluate the plausibility of these features with a detailed interpretation of these features in a biological context.

We therefore elucidate to what extent the top features are biologically plausible in the context of AD, and whether this offers a possible explanation for the improved classification performance with simulated features.

Interestingly there is some regional overlap between the top features related to Aβ and Tau, namely the left entorhinal cortex and the right fundus of the superior temporal sulcus. This overlap could represent a potential synergistic effect between Aβ and Tau in these regions. Synergistic effects between Aβ and Tau in the temporal lobe are an important element in the progression from MCI to AD. They have been described in voxel-based analyses, as well as in molecular studies [3-5]. In a study by Halawa, Gatchel [6] both temporal tau and amyloid burden were associated with an impairment of activities of daily living. Still, the combination of both pathologic markers showed an association that by far surpassed the mere additive effect.

High-ranked features in the visuospatial system may correspond to alterations of memory processes in the visuospatial and episodic compartment and were also used to classify between MCI and HC [7, 8].

Higher Aβ-burden in older adults was associated with decreased neural activity in the ventral visual stream [9] and Aβ-PET positive MCI patients perform worse in a navigation task than Aβ-PET negative MCI patients [10]. Other essential contributors to spatial navigation in the brain’s navigation circuit are the grid cells in the entorhinal cortex, which is primarily affected by tau deposition [11]. Tau-related disruption of grid-cell firing in the entorhinal cortex of transgenic mice expressing human tau leads to increased theta oscillations that were associated with spatial memory deficits [12]. In humans, the performance in an entorhinal cortex-based immersive virtual reality navigation task helped to differentiate between MCI patients at low and high risk of developing AD [13].

On a network level, theta-band oscillations in high-density EEG in a network consisting of the temporal lobe, striatum, inferior occipital lobe, and cerebellum correlated with the performance in a spatial memory task [14] and the reduction in visual processing network complexity correlated with the stage of AD [15]. Interestingly, a recent review focused on spatial navigation measured with LFPs revealed that the neural representation of spatial features occurs on a mesoscopic level of mainly theta oscillations [16] and thus the same level that TVB operates [17].

The default mode network (DMN) was exclusively represented by empirical top features. The DMN is consistently affected in AD [18] and shows the earliest accumulation of Aβ [19]. Nevertheless, the empirical features poorly performed when classifying MCI, which can in some cases be interpreted as a prodromal stage of AD [20]. This could partly be explained by the missing network information of the empirical features. The breakdown of functional connectivity of the DMN occurs concurrently with Aβ deposition in preclinical subjects [21] and in the early stage of MCI [22]. It is possible that network disturbances, detectable by the simulated features, occur even earlier than Aβ deposition in the DMN, but simulated features do not represent the DMN. Alves et al. [23] proposed a new neuroanatomical model of the DMN that includes subcortical structures like the thalamus, which is represented by the simulated top features. Following this approach, the combination of empirical and simulated top features yields a more comprehensive description of this functional network that is an important marker in the disease progression of AD.

Network disturbances (as seen in the subcortical hub regions) may come apparent earlier than stronger Aβ or Tau uptake in the PET-Imaging leading to a better classification of the MCI group. At the same time, simulated features only use the amyloid PET and cannot get the information from the Tau-PET, which seems to better correlate with the cognitive decline of patients [24]. This missing information could therefore be a reason for the misclassification of HC by the simulated features. Hence, it would be of great interest how a simulation using Tau-PET performs in this classification problem.

Finally, we observed high importance of thalamic frequencies. In recent years there has been considerable evidence that thalamic dysfunction. Mainly the anterior thalamus has been identified to play a crucial role in early disease progression [25]. The anterior thalamus is part of the Papez circuit and as such strongly connected to episodic memory [26]. While the relevance of the (unparcellated) thalamus is represented by the simulated features, it did not play an important role as an empirical feature. This disparity makes sense for Tau, considering that only some of the thalamic nuclei have been shown to contain NFT even in late stages of the disease [27] and therefore, a strong PET-signal is unlikely. In MCI patients, the thalamus displays both increased and decreased functional connectivity, which could be a hint towards loss of function or compensation mechanisms [28]. A change in the LFP of the thalamus supports a shift in function, as different frequency bands can be attributed to various brain functions [29]. According to Schnitzler and Gross [30], faster rhythmic activities correspond to more local neuronal communication, while slower oscillations likely arise from larger populations within wider-range networks. A change in the function of the thalamus and other subcortical hubs could therefore have possible network effects detectable in LFP/EEG. Reduced complexity of the EEG signal and perturbations in EEG synchrony are significant effects of AD on EEG [31, 32].
